# Temporary parasitic ants modify nestmate discrimination patterns of host workers

**DOI:** 10.1101/2025.05.10.653236

**Authors:** Ryotaro Kobayashi, Yasukazu Okada, Toshiharu Akino, Naoto Idogawa

**Author notes:** Corresponding authors (R. Kobayashi), Laboratory of Animal Sociology, Department of Biology, Graduate School of Science, Osaka Metropolitan University, 3-3-138 Sugimoto, Sumiyoshi, Osaka 558-8585, Japan, (N. Idogawa), Group of Integrative Ecology and Evolution, Department of Biological Science, Nagoya University, Furoucho, Chikusa-ku, Nagoya 464-8602, Japan.

## Abstract

Social insects rely on nestmate discrimination systems to maintain highly organized societies. Some social parasites can evade these systems to exploit their hosts. In this study, we report that the temporarily socially parasitic ant *Lasius umbratus* may modify the nestmate discrimination patterns of its host, *L. japonicus*. Aggression assays revealed that unparasitized hosts exhibited high aggression toward non-nestmate conspecifics and parasites. In contrast, parasitized hosts showed significantly lower aggression toward non-nestmate conspecifics and parasites, while maintaining considerable aggression toward a non-parasitic congener as a positive control. This suggests that parasites modify the nestmate discrimination patterns of the hosts rather than merely suppressing aggression.

Furthermore, chemical analysis of cuticular hydrocarbons revealed that hosts from mixed-species colonies of hosts and parasites had profiles resembling those of unparasitized hosts, whereas parasites from mixed-species and parasite-only colonies showed distinct profiles. Notably, parasites from mixed-species colonies had hydrocarbon profiles resembling those of their hosts, indicating that *L. umbratus* employs chemical mimicry to alter host behaviour. Our findings shed light on the intricate strategies of social parasites and the behavioural and chemical mechanisms sustaining highly organized ant societies.

## INTRODUCTION

In group-living animals, such as flocks or familial groups, the elimination of intruders is essential because intrusion incurs significant costs, including a decrease in the number of descendants. (Emlen and Wrege 1986; Blažek et al. 2018; Buschinger 2009). Various cues are used by animals to recognize and differentiate between individuals, such as auditory signals in birds (Colombelli-Négrel et al. 2012; Thorpe 1958) and olfactory and visual cues in fish and mammals (Brennan and Kendrick 2006; Satoh et al. 2016; Ward et al. 2020). This discrimination between self and others, along with several sensory modalities, functions as a social immunity system, expelling intruders and preserving group integrity (Cotter and Kilner 2010; Cremer et al. 2007).

However, some parasites can evade host discrimination systems. For example, cuckoos employ visual deception by laying mimicking eggs in the nests of their host species (Langmore et al. 2003). Furthermore, cuckoo chicks engage in auditory mimicry by producing calls similar to those of host chicks, thereby exploiting parental care (Langmore et al. 2003). Cuckoo catfish disrupt the spawning behaviour of the host to lay eggs (Sato 1986), which are then cared for by the host alongside its own eggs (Blažek et al. 2018). Finally, the hatched fry consumes the eggs of the host. Such brood parasitism occurs across a broad range of taxa (Summers and Amos 1997; Tallamy 2005; Wisenden 1999), offering valuable insights into the mechanisms for eliminating intruders and revealing the vulnerabilities underlying social systems in animals.

Brood parasitism has evolved multiple times in insects (Sless et al. 2023). It is referred to as “social parasitism” in social Hymenoptera and has been reported in at least 492 species (Rabeling 2021). Social Hymenoptera defend against social parasites by distinguishing nestmates through colony-specific cuticular hydrocarbon (CHCs) profiles (Akino et al. 2004; van Zweden and d’Ettorre 2010). They reject non-nestmates, even of the same species (Bos and d’Ettorre 2012; Martin et al. 2008; van Zweden and d’Ettorre 2010; Villalta et al. 2020). Thus, social parasites must deceive the chemical nestmate discrimination system of the hosts to coexist and exploit their resources (Hölldobler and Wilson 1990; Lenoir et al. 2001; Rabeling 2021).

Parasites coexist with their hosts by employing three chemical mimicry strategies, as outlined below: 1) mimicking host odour: parasites replicate the host’s odour either by grooming the host (Lenoir et al. 2001) or through the biosynthesis of odour chemicals (Bauer et al. 2010; Dettner and Liepert 1994; Howard et al. 1982); 2) modifying the odour of the host colony: parasites introduce characteristic odours, thereby altering the odour of the colony (Bagnères et al. 1996; Lorenzi 2006; Turillazzi et al. 2000); and 3) avoiding detection by the host: parasites employ cuticular odours that remain either undetected (Jeral et al. 1997) or insignificant (Cervo et al. 2008) to the host, thereby evading discrimination and subsequent attacks (Lenoir et al. 2001; Lorenzi and d’Ettorre 2020; Neupert et al. 2018).

Social parasitism involves interactions between the parasites and their hosts. Therefore, for a comprehensive understanding, it is essential to focus not only on parasite signalling, such as chemical mimicry, but also on changes in the nestmate recognition patterns and decision-making processes of the hosts after they receive the signals. In this study, we aimed to investigate how host-parasite interactions change with social context by examining the behavior of the ant *Lasius japonicus* and its temporary social parasite *L. umbratus* (Quque and Bles 2021; Seifert 1992). To elucidate the nestmate discrimination patterns of hosts, we compared the aggressive responses of parasitized and unparasitized hosts toward different types of opponents, including nestmates, non-nestmates, congeners, conspecifics, and parasites. Additionally, we analyzed the cuticular hydrocarbons of both the host and parasite in the monospecific and mixed colonies to examine the chemical basis of the parasitic strategy.

## METHODS

### Materials

Host species *L. japonicus*, parasite species *L. umbratus*, and their congener *L. hayashi* were maintained in the laboratory for the experiments described in later sections. Experimental colonies of the host and parasite species were established using newly mated queens captured immediately after their nuptial flights. Foundress queens of *L. japonicus* were collected in June 2022 from Minamiosawa, Hachioji City, Japan (latitude: 35.620557° N, longitude: 139.381205° E), and each was reared separately in the laboratory to found the colonies. Mated queens of *L. umbratus*, collected in June 2022 from the same location, were introduced into artificial colonies of *L. japonicus*, comprising 50 workers and were maintained for approximately one year. Workers of *L. hayashi* were collected in May 2023 from Takao, Hachioji City (latitude: 35.624210° N, longitude: 139.244586° E), and in September 2023 from Minamiosawa, Hachioji City, and subsequently reared.

All colonies were kept in plastic containers (H × D × W = 60 × 189 × 130 mm) maintained at a temperature range of 25–27 °C with a light: dark cycle of 12:12 h. Each colony was provided with an artificial nest constructed from plastic Petri dishes, plaster, and transparent red PVC sheets. The ants were fed insect jelly (Pro jelly, Oreos Ltd., Saitama Prefecture, Japan) and frozen crickets (House crickets, Tsukiyono-farm Ltd., Gunma Prefecture, Japan) every three days in sufficient quantities.

### Aggression assay

We examined changes in the patterns of nestmate discrimination between parasitized and unparasitized host workers by comparing their aggressive behavior toward various types of opponents.

The aggressive response was assessed using a 1 vs. 1 arena assay of forager workers. In the assay, a focal worker and an opponent worker were gently transferred to a circular arena (plastic vials, inside diameter= 15 mm, the wall surface was coated by Fluon® to obstruct climbing by ants). The behaviour of the focal worker was observed and recorded for 5 min using a video camera (Logicool HD Pro Webcam C920). We scored the aggressive behaviour of a focal worker toward an opponent for 5 min using the following categories in order of escalating aggression scores, as modified from previous studies (Steiner et al. 2007; Suarez et al. 2002): 0 = ignore (no contact between both workers or no response, even if contact occurs), 1 = prolonged antennation, 2 = avoiding opponent worker, 3 = aggression (focal worker lunging at the opposing ant), and 4 = fight (prolonged aggression and biting). At the end of 5 min, the maximum score during the trial was recorded. Each worker participated in only one trial.

Foragers were defined as individuals who were observed foraging outside the nest during feeding. They were identified and marked on their dorsal abdomen using a paint marker (Paint Marker PX206C, Mitsubishi Pencil Company Ltd., Tokyo, Japan) at least 12 h prior to the trial.

### Opponent pairings of aggression assays

We measured the aggression levels of unparasitized and parasitized *L. japonicus* workers under ten experimental conditions (Table 1). In the experimental design, focal unparasitized host workers were paired with either conspecific non-nestmates (T6) or non-nestmate parasitic *L. umbratus* (T4). Focal parasitized host workers were paired with conspecific non-nestmates (T7), nestmate parasites (T3), or non-nestmate parasites (T5). For the negative control, focal unparasitized hosts (T1) and focal parasitized hosts (T2) were paired with their conspecific nestmates. As the positive control, the non-parasitic congener *L. hayashi* (Boudinot et al. 2022) was introduced into both focal unparasitized hosts (T8) and focal parasitized hosts (T9). To examine the effects of queen absence on aggression, orphan colonies of the host *L. japonicus* were created by removing the queen from unparasitized colonies. The focal orphan colonies were paired with their conspecific non-nestmates (T10).

**Table 1.** Conditions of opponent pairings for aggression assays.

| Test ID | Focal host workers (*L. japonicus*) | Opponent worker's conditions |
| --- | --- | --- |
| T 1 | Unparasitized | conspecific nestmate *L. japonicus* |
| T 2 | Parasitized | conspecific nestmate *L. japonicus* |
| T 3 | Parasitized | nestmate parasite *L. umbratus* |
| T 4 | Unparasitized | non-nestmate parasite *L. umbratus* |
| T 5 | Parasitized | non-nestmate parasite *L. umbratus* |
| T 6 | Unparasitized | conspecific non-nestmate *L. japonicus* |
| T 7 | Parasitized | conspecific non-nestmate *L. japonicus* |
| T 8 | Unparasitized | non-nestmate *L. hayashi* |
| T 9 | Parasitized | non-nestmate *L. hayashi* |
| T 10 | Orphan | conspecific non-nestmate *L. japonicus* |

Each condition was replicated 25 times, with colonies randomly selected from multiple identical rearing conditions for each trial (Supplemental file 1). All behavioural data from our findings are shown in Supplemental file 2.

### Comparing aggression

Aggression levels across all conditions were compared pairwise using the Wilcoxon rank-sum test (adjusted by the Holm–Bonferroni method). Statistical analyses were conducted using R version 4.2.3 (R Core Team, 2023).

### Chemical Analysis

To investigate chemical mimicry during parasitism, we compared the cuticular hydrocarbon profiles of parasitic *Lasius umbratus* and its host, *Lasius japonicus*, under four specific conditions: Js (*L. japonicus* worker from monospecific colonies); Us (*L. umbratus* worker from monospecific colonies); Jm (*L. japonicus* worker from mixed-species colonies); and Um (*L. umbratus* worker from mixed-species colonies).

For each condition, four randomly selected foragers from each of the three replicate colonies were killed by freezing for at least 10 min. Individual workers were immersed in 10 μL of hexane for 3 min, and 2 μL of the extracts were analyzed using gas chromatography (GC). GC analyses were performed on the gas chromatograph Shimadzu GC-2014 equipped with a flame ionization detector and a non-polar column GL Sciences InertCap1 (15 m length, 0.25 mm inside diameter, 0.25 μm film thickness). The injection was performed in splitless mode for 1 min, and the flame ionization detector was set at 300 °C. The oven temperature was held at 40 ºC for 5 min, programmed to increase from 40 ºC to 300 ºC at 20 ºC/min, and maintained at 300 ºC for 10 min. The column head pressure was set to 40 kPa, and helium was used as the carrier gas. The GC data were processed using a Shimadzu Chromatopak CR8A instrument. All GC data are provided in Supplemental file 3.

### Principal component analysis of cuticular hydrocarbons

As species specific peaks were not obvious from the chromatograms (Supplemental file 4), we selected 13 peaks in the C28 to C34 which were detected in both *L. japonicus*, and *L. umbratus*s, and these were used to compare the chemical profiles across all experimental conditions. All peaks were normalized, and the area percentages were input into a principal component analysis (PCA) matrix. The significance of the component differences was determined via permutational multivariate analysis of variance (PERMANOVA) with Bray-Curtis similarities and 1000 permutations. In the PERMANOVA, a single factor with four levels was used, corresponding to the following groups: monospecific *L. japonicus* colonies (Js), monospecific *L. umbratus* colonies (Us), *L. japonicus* from mixed-species colonies (Jm), and *L. umbratus* from mixed-species colonies (Um). PCA and PERMANOVA were performed using R ver. 4.2.3(R Core Team, 2023).

### Physiological status of unparasitized and parasitized hosts

The fat content and ovarian development of unparasitized and parasitized host (*Lasius japonicus*) workers were analyzed to assess the physiological status of nurse workers. Four separate unparasitized host colonies were divided into four unparasitized and four parasitized colonies. Each colony contained approximately 120 workers and 30 mg eggs or larvae. All colonies were maintained under the same breeding conditions as previously described. Worker fat content and ovarian development were assessed for approximately two months (62–71 days, n = 9) after colony establishment.

Nurse workers are defined as individuals who care for broods within a nest during feeding.

### Measurement of fat content

Two months after the colony establishment, the fat content of the workers was measured. Ten workers from each colony were assessed for their wet weight and head width. Subsequently, the workers were subjected to fat content analysis using the methanol-chloroform method (Barnes and Blackstock 1973; Idogawa et al. 2017; Kishino et al. 2024). The workers were then dried at 65 °C for 24 hours, and their dry mass was determined to the nearest 0.01 mg using an MS105 ultra-microbalance (Mettler-Toledo, Switzerland). Subsequently, fat was extracted using a solvent composed of a 2:1 mixture of chloroform and methanol (Wako, Japan) by immersing the dried samples in the solvent (0.5 mL) for 2 days at room temperature. Following fat extraction, ants were dried again at 65 °C for 25 h and weighed again to determine their lean mass. Body fat content was calculated as the body fat percentage using the formula: (dry mass□− □lean mass) □× □100/dry mass. Fat content data was analyzed by linear mixed models (LMM) in the lmer function of the “lme4” package (Bates et al. 2015). Individuals were scaled by head width and wet weight and colony ID was applied as a random factor. All fat content data from this study are shown in Supplemental file 6.

### Ovary dissection and evaluation of reproductive state

Two months after colony establishment, the fat content of the workers was analyzed. Ten workers from each colony were measured for wet weight and head width. Subsequently, the ovarian state of each individual was examined by dissection. Dissections were performed in 1× PBS and individual ovaries were imaged using a stereomicroscope (S9i, Leica, Germany). Ovarian activation scores were categorized into four stages based on previous studies (Holman et al. 2010); (1) completely empty; (2) one or two very small eggs and/or developing nurse cell material; (3) one to three developing eggs in both ovarioles or large eggs in one ovariole; and (4) well-developed eggs in both ovarioles. Ovarian activation scores were ordinal dependent variables, so we applied an ordered logistic error distribution in the CLMM function of the “ordinal” package (Christensen 2024). Individuals were scaled by head width and wet weight and colony ID was applied as a random factor. All ovarian activation data from our study are shown in Supplemental file 7.

### Ethical note

No licenses or permits were required for this research, as *Lasius japonicus, L. umbratus*, and *L. hayashi* are widely distributed and not protected species in Japan. Individuals of each species were carefully collected from areas with abundant local populations, and only the minimum number of individuals necessary to meet the statistical requirements of the study was used. To minimize stress caused by artificial conditions, all ants were housed in humidified plaster nests equipped with foraging areas. Colonies were maintained at optimal temperatures and were continuously supplied with sufficient food and water to reflect near-natural conditions. In behavioural experiments, individuals were, due to experimental design, isolated from their original groups. However, since *Lasius* ants cannot be maintained in good health when housed alone, those used in behavioural assays were humanely euthanized by freezing at −20□°C immediately after the experiments. Ants used for chemical analyses and Comparison of physiological status were euthanized in the same manner prior to analyses.

## RESULTS

*Aggression assay*

The aggression assay revealed that both unparasitized (T1) and parasitized (T2) hosts exhibited low levels of aggression toward conspecific nestmates, as observed in the negative control (Wilcoxon rank sum test, Holm-adjusted: *W* = 325, *P* = 1) (Fig. 1; T1, T2). Similarly, parasitized hosts exhibited low levels of aggression toward the parasitic nestmate *L. umbratus* (T3), with no significant difference compared to aggression levels toward conspecific nestmates (T2) (Wilcoxon rank-sum test, Holm-adjusted: *W* = 300, *P*= 1) (Fig. 1; T3, T2). Unparasitized hosts exhibited significantly higher aggression toward non-nestmate conspecifics (T6) and parasites (T4) than toward conspecific nestmates (T1), (Wilcoxon rank-sum test, Holm-adjusted: *W* = 53, *P* < 0.001) (Fig. 1; T6, T1), (Wilcoxon rank-sum test, Holm-adjusted: *W* = 53, *P* < 0.001) (Fig. 1; T4, T1). In contrast, parasitized hosts displayed low levels of aggression toward non-nestmates, regardless of whether they were conspecifics (T7) or parasites (T5). These levels were comparable to the levels observed toward conspecific nestmates (T2), (Wilcoxon rank-sum test, Holm-adjusted: *W* = 237.5, *P*= 0.187) (Fig. 1; T7, T2), (Wilcoxon rank-sum test, Holm-adjusted: *W* = 237.5, *P* = 0.187) (Fig. 1; T5, T2).

**Figure 1.**
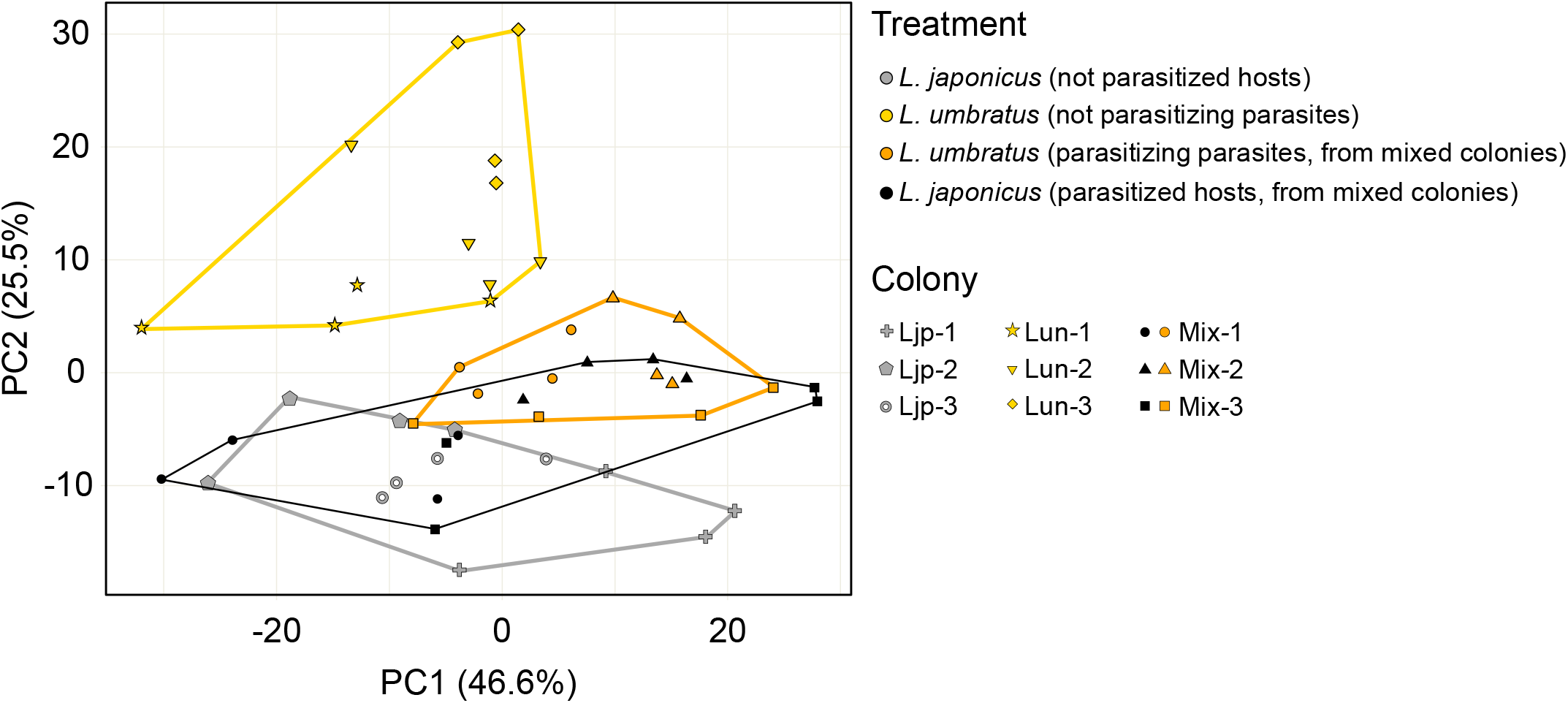
Aggression levels of unparasitized (u), parasitized (p), and orphan (o) host workers. Aggression scores reflect the behavior of the focal host workers: 0 = no response, 1 = touch, 2 = avoid, 3 = aggression, and 4 = fight. A total of 25 replicates were included for each condition. Circle sizes represent the number of observations for each aggression score. The letters at the top of the figure indicate significant differences in aggression levels between conditions (Wilcoxon rank-sum test, Holm-adjusted: *P* < 0.05).

An assay measuring aggression toward the non-parasitic congener *L. hayashi* as a positive control revealed that both unparasitized (T8) and parasitized (T9) hosts exhibited high levels of aggression (Wilcoxon rank-sum test, Holm-adjusted: *W* = 313, *P* = 1) (Fig. 1; T8, T9).

Moreover, host workers from orphan colonies exhibited high levels of aggression toward conspecific non-nestmates (T10). The aggression levels were not significantly different from those of the unparasitized hosts (T6) (Wilcoxon rank sum test, Holm-adjusted: *W* = 209, *P* = 0.37) (Fig. 1; T10, T6), but were significantly higher than those of the parasitized hosts (T7) (Wilcoxon rank-sum test, Holm-adjusted: *W* = 465.5, *P*= 0.03) (Fig. 1; T7, T10). All statistical analysis data are presented in Supplemental file 5.

### Analysis of cuticular hydrocarbons

Principal component analysis (PCA) of the 13 peaks detected in both *L. japonicus* (host) and *L. umbratus* (parasite) workers demonstrated that principal components 1 (accounting for 46.6% of the variance) and 2 (accounting for 25.5% of the variance) collectively accounted for 72% of the total variance. Significant differences in cuticular hydrocarbon profiles were identified between *L. japonicus* workers from monospecific colonies (Js) and *L. umbratus* workers from monospecific colonies (Us) (PERMANOVA: *P* = 0.0006, Fig. 2), between *L. japonicus* workers from monospecific colonies (Js) and *L. umbratus* workers from mixed-species colonies (Um) (PERMANOVA: *P* = 0.0048, Fig. 2), between *L. umbratus* workers from monospecific colonies (Us) and *L. umbratus* workers from mixed-species colonies (Um) (PERMANOVA: *P* = 0.0006, Fig. 2), and between *L. japonicus* workers from mixed-species colonies (Jm) and *L. umbratus* workers from monospecific colonies (Us) (PERMANOVA: *P* = 0.0006, Fig. 2).

**Figure 2.**
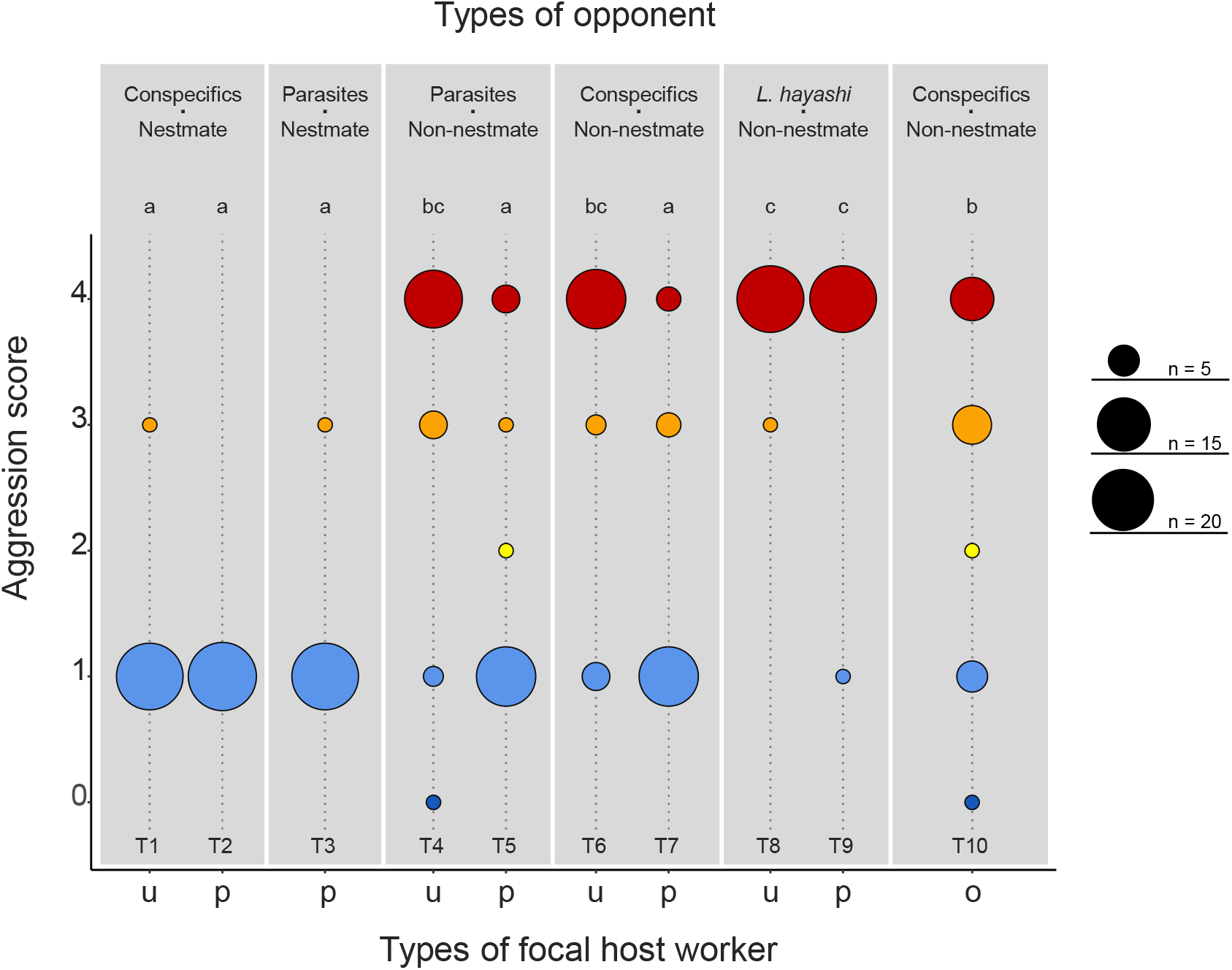
Principal Component Analysis (PCA) plot based on gas chromatogram data from four treatments. Colors indicate treatment groups, with each group enclosed within a convex hull. Shapes denote the source colonies.

In contrast, no significant differences were observed between *L. japonicus* workers from monospecific colonies (Js) and *L. japonicus* workers from mixed-species colonies (Jm) (PERMANOVA: *P* = 0.5423, Fig. 2) or between *L. japonicus* workers from mixed-species colonies (Jm) and *L. umbratus* workers from mixed-species colonies (Um) (PERMANOVA: *P* = 0.6485, Fig. 2).

### Physiological status of unparasitized and parasitized hosts

The fat content was not significantly different between the unparasitized (mean ± SD□= □32.9 ± 12.7 %) and parasitized colonies (mean□± □SD□=L34.4□± □13.1 %) (LMM, estimate = -0.0133, *SE* = 0.0165, *df* = 85.6, *t*-ratio = -0.809, *P* = 0.4210) (Supplemental file 1). Similarly, ovarian development was not significantly different between unparasitized and parasitized colonies (CLMM, estimate = 0.0599, *SE* = 0.381, *df* = Inf., *z*-ratio = 0.157, *P* = 0.8752) (Supplemental file 1).

## DISCUSSION

In the present study, we focused on nestmate recognition by hosts to clarify the mechanisms by which temporary social parasites coexist with their hosts. Behavioural observations confirmed that parasitized *L. japonicus* workers exhibited very low levels of aggression toward the nestmate parasite *L. umbratus* workers were very low (Fig. 1; T3), showing no significant difference from aggression levels toward conspecific nestmates (Fig. 1; T2). In contrast, unparasitized host workers exhibited significantly higher levels of aggression toward *L. umbratus* (Fig. 1; T4), suggesting that *L. umbratus* workers were not unconditionally accepted by *L. japonicus*. Interestingly, parasitized host workers exhibited low levels of aggression toward parasites (Fig. 1; T5) and conspecific non-nestmates (Fig. 1; T7). However, unparasitized hosts typically attacked and eliminated non-nestmate conspecifics (Fig. 1; T6). Together with the high levels of aggression toward the non-parasitic congener *L. hayashi* (Boudinot et al. 2022) workers observed in both parasitized and unparasitized *L. japonicus* workers (Fig. 1; T8, 9), our results suggest that the presence of parasitic *L. umbratus* does not suppress attack behaviour itself but rather modifies the nestmate discrimination patterns of the host *L. japonicus* workers.

One possible socioenvironmental factor that influences the aggression level of host workers is the absence of a host queen. Several studies on social insects have reported a decrease in worker aggressiveness after the death of the foundress queen (Stuart and Herbers 2000; Vander Meer and Alonso 2002). In our experiment, the parasitized *L. japonicus* colonies did not include their mother queen but included the parasitic *L. umbratus* queen. To examine the influence of the absence of a conspecific foundress queen, we removed the queen from the unparasitized colonies and measured worker aggression levels against non-nestmate conspecifics. There was no significant difference in aggressiveness between the orphaned group (Fig. 1; T10) and the queenright group (Fig. 1; T6). In contrast, parasitized workers showed significantly lower aggression levels than orphaned workers (Fig. 1, T7). Thus, the absence of the original queen does not seem to be the primary cause of the reduced aggression observed in the parasitized hosts. Furthermore, comparisons of fat content and ovarian development between nurse workers from unparasitized and parasitized colonies revealed no significant differences (Supplemental file 1). Therefore, it is unlikely that social parasitism imposes physiological stress on the host, resulting in changes in its behavioural patterns.

Social parasites evade host attacks by employing chemical mimicry (Lenoir et al. 2001). This strategy involves mimicking the cuticular hydrocarbon profiles of the host colony, which are highly colony-specific and essential for nestmate recognition in Hymenoptera (Turgis and Ordon 2012; van Zweden and d’Ettorre 2010). Chemical mimicry can be categorized into three main strategies: 1) mimicking the host’s odour (Bauer et al. 2009; Howard et al. 1982), 2) altering the odour of the host colony (Turillazzi et al. 2000), and 3) evading detection by the host (Jeral et al. 1997; Lambardi et al. 2007). Additional chemical-based strategies for avoiding attacks include emitting repellent odours (Lhomme et al. 2012) or releasing chemicals that suppress aggression (Elia et al. 2018; Martin et al. 2007).

In the present study, gas chromatographic analysis revealed that workers from monospecific colony of *L. umbratus* exhibited cuticular hydrocarbon (CHC) profiles that were distinct from those of *L. japonicus* (Fig. 2). However, parasitic *L. umbratus* workers showed CHC compositions similar to those of their host, *L. japonicus* (Fig. 2), supporting the first strategy of chemical mimicry via host odour imitation (Bauer et al. 2009; Howard et al. 1982), where the parasite imitates the CHC composition of the hosts. This alteration in CHC composition is consistent with the results of aggression assays, indicating that *L. umbratus* workers were not attacked by their hosts. However, the observation that even non-nestmate *L. japonicus* workers, which do not employ chemical mimicry, are not attacked by parasitized *L. japonicus* workers requires a different explanation.

We propose that in addition to mimicking the CHC composition of the host, *L. umbratus* may alter the recognition mechanisms of its host, *L. japonicus*. Changes in nestmate recognition patterns are known in other social parasitism systems, where the parasite overlays its own CHC on the host. For example, in parasitized colonies of the socially parasitic wasp *Polistes atrimandibularis*, host aggression toward nestmates increases, whereas aggression toward non-nestmates decreases, likely due to errors in nestmate recognition caused by the mixing of chemical signatures (Lorenzi 2003). In contrast, *L. japonicus* colonies exhibited no changes in the CHC profiles of individual members, whether parasitized or not, suggesting that behavioural changes are driven by modifications in internal recognition processes rather than alterations in chemical signatures.

Although the specific mechanisms underlying these changes remain unclear, transcriptomic analyses of the brains and antennae of parasitized hosts in other obligate social parasitic species, such as *Harpagoxenus sublaevis*, have revealed differences in the expression of genes related to olfactory perception compared to non-parasitized hosts (Stoldt et al. 2023). Such neurobiological investigations are warranted in *L. japonicus* to better understand the neurophysiological processes involved in social parasitism. Moreover, other socially parasitic species may exhibit comparable alterations in host recognition patterns. Further studies are essential to elucidate the generality of such mechanisms in social parasitism systems.

This study provides a novel perspective that during the process of being accepted by hosts, social parasites may not only employ chemical mimicry but also modify the nestmate discrimination systems of the hosts. Further investigation into the cognitive mechanisms and behavioural physiology of hosts in the context of social parasitism will deepen our understanding of social parasitism and provide valuable insights into the mechanisms that sustain insect societies.

## Supporting information

Table presenting the number colonies used as focal and opponent, and figure of comparison of the physiological status of unparasitized and parasitized

All aggressive behavioural data of focal hosts Lasius japonicus support our findings.

Cuticular hydrocarbons data by GC analyses supporting our findings in this study.

Typical examples of gas chromatogram for each species and conditions.

Detailed statistical analysis results by Wilcoxon rank sum test.

All fat contents data for comparison of physiological status of host ants.

All ovarian activation data for comparison of physiological status of host ants.

## Author Contributions

**Ryotaro Kobayashi**: data curation, investigation, methodology, visualization, writing—original draft and writing—review and editing. **Yasukazu Okada**: project administration, supervision, data curation, methodology, funding acquisition, writing—review and editing. **Toshiharu Akino**: investigation, methodology, resources. **Naoto Idogawa**: investigation, conceptualization, supervision, methodology, visualization, funding acquisition, writing—review and editing, funding acquisition, formal analysis. All authors gave final approval for publication and have agreed to be held accountable for the work performed therein.

## Data Availability

Data used for the analyses described here can be found in the Supplementary material.

## Acknowledgements

We thank T. Wakamiya, K. Kishino, K. Kawamoto, and members of the Animal Ecology Lab at Tokyo Metropolitan University for fruitful discussions. We would like to thank Y. Mitaka for valuable advice regarding chemical analysis. This work was supported by the Japan Society for the Promotion of Science (JSPS) Research Fellowship for Young Scientists to NI (22KJ2563) and a grant for young scientists from Tokyo Metropolitan University to YO. The founders played no role in the study design, data collection, and analysis, decision to publish, or manuscript preparation.

## Supplementary material

**Supplemental file 1**

Table presenting the number colonies used as focal and opponent, and figure of comparison of the physiological status of unparasitized and parasitized hosts.

**Supplemental file 2**

All aggressive behavioural data of focal hosts *Lasius japonicus* support our findings.

**Supplemental file 3**

Cuticular hydrocarbons data by GC analyses supporting our findings in this study.

**Supplemental file 4**

Typical examples of gas chromatogram for each species and conditions.

**Supplemental file 5**

Detailed statistical analysis results by Wilcoxon rank sum test.

**Supplemental file 6**

All fat contents data for comparison of physiological status of host ants.

**Supplemental file 7**

All ovarian activation data for comparison of physiological status of host ants.

