## Supplementary material for "Temporary parasitic ants modify nestmate discrimination patterns of host workers": Table presenting the number colonies used as focal and opponent, and figure of comparison of the physiological status of unparasitized and parasitized

**Supplemental Materials**

Supplemental table S1: The experimental design and the number of colonies used as focal and opponent

| **Test ID** | **The number of focal colonies** | **The number of opponent colonies** |
| --- | --- | --- |
| T 1 | 10 | 10 |
| T 2 | 7 | 10 |
| T 3 | 4 | 4 |
| T 4 | 10 | 4 |
| T 5 | 4 | 3 |
| T 6 | 5 | 8 |
| T 7 | 5 | 10 |
| T 8 | 10 | 1 |
| T 9 | 4 | 1 |
| T 10 | 4 | 10 |

**Comparison of** **physiological status of unparasitized and parasitized hosts**

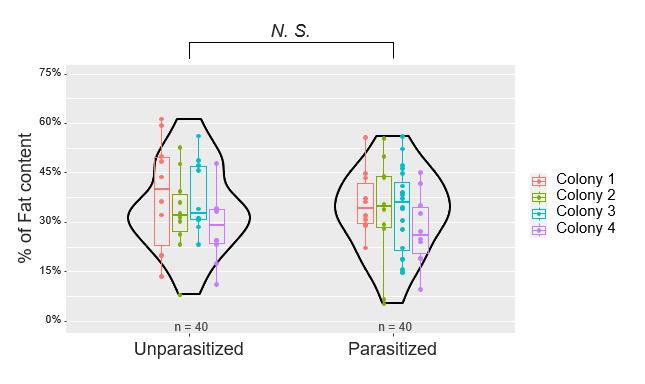

Supplemental figure S1.	Fat content of unparasitized and parasitized host nurse workers measured two months after colony establishment (LMM, estimate = -0.0133, SE = 0.0165, *df* = 85.6, t-ratio = -0.809, *p* = 0.4210).

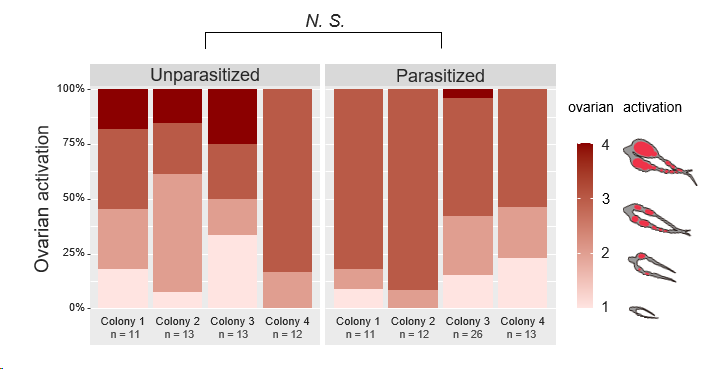

Supplemental figure S2:	Ovarian activation in unparasitized and parasitized host nurse workers measured two months after colony establishment (CLMM, estimate = 0.0599, SE = 0.381, *df* = *Inf.*, z-ratio = 0.157, *p* = 0.8752).
