## Supplementary figures and images for "Temporary parasitic ants modify nestmate discrimination patterns of host workers"

### Typical examples of gas chromatogram for each species and conditions.

#### Typical examples of gas chromatogram for each species and conditions

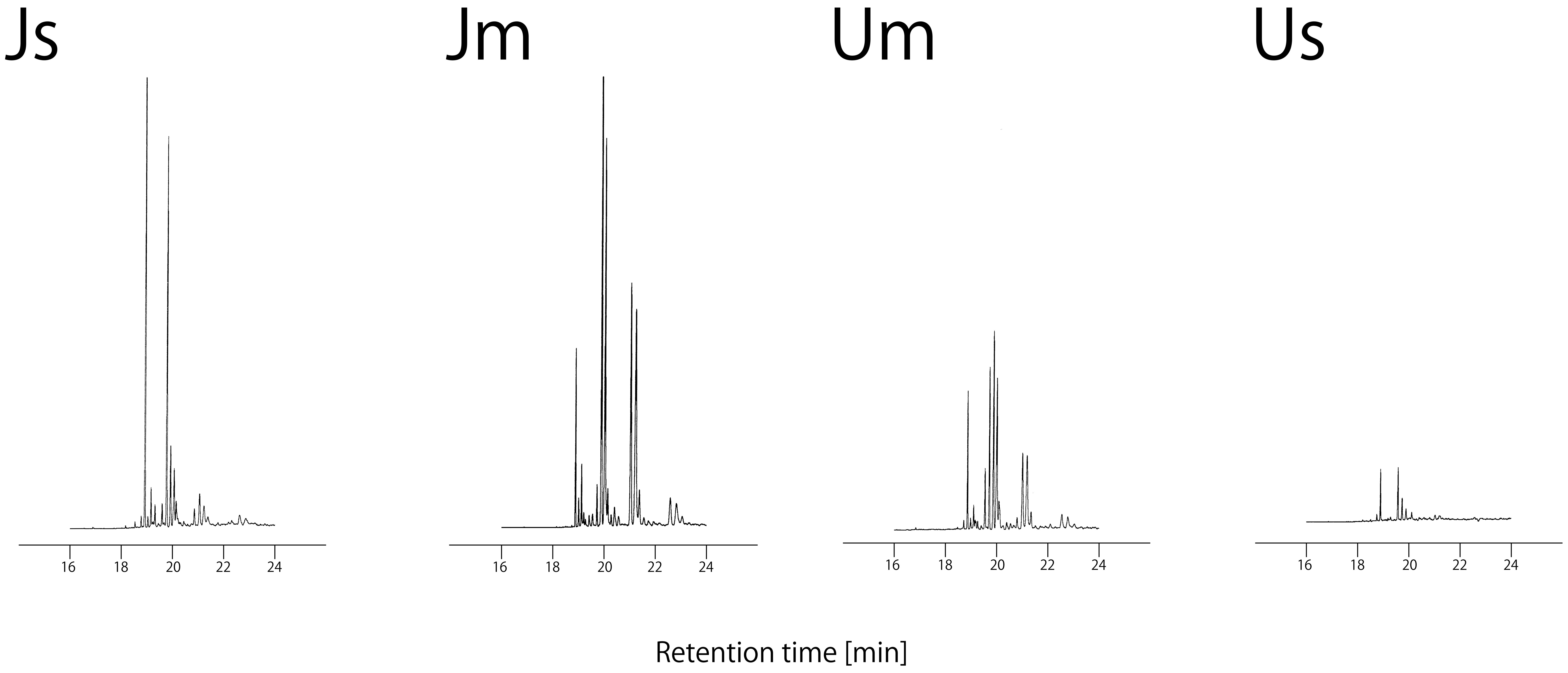
